# Opioid withdrawal engages a habenular subpopulation responsive to aversive states

**DOI:** 10.64898/2026.04.29.721196

**Authors:** Leah G Blankenship, Vincent Moore, Blake Holcomb, Isabella Salinas, Emily L Sylwestrak

## Abstract

Opioid withdrawal produces a protracted aversive state that is a driving factor towards relapse for patients with opioid use disorder. The habenula is known to mediate symptoms of withdrawal across several substance use disorders, including alcohol, nicotine, and opioid use disorder. However, it is unclear which habenular populations contribute to withdrawal symptoms. Here, we identify a cell type marker for habenular neurons active during opioid withdrawal. Using immediate early gene analysis, we find that glutamic acid decarboxylase 2-expressing (GAD2^+^) cells in the lateral habenula (LHb^GAD2^) increase activity in response to both spontaneous and naloxone-precipitated opioid withdrawal in mice. Recording neural activity *in vivo* revealed transient activity at the presentation of aversive stimuli, and a sustained increase in activity during aversive states such as opioid withdrawal and inescapable shock. These results highlight the cell type heterogeneity of the habenula, and the role of specific cell types in opioid withdrawal.

## Introduction

Deaths from opioid overdose rose 64% from 2019 to 2022 in the United States, reaching 81,806 deaths in 2022.^1^ Addiction is a self-perpetuating cycle: an individual binges the substance of abuse, endures withdrawal symptoms during abstinence, and craving during withdrawal leads to binging, starting the cycle anew.^2–4^ The desire to alleviate withdrawal symptoms is an important factor driving relapse, and overdose is more likely during relapse due to reduced tolerance.^2–5^ Thus, preventing or mitigating opioid withdrawal symptoms is critical for reducing harm and mortality in patients with opioid use disorder. Somatic symptoms of opioid withdrawal include muscle pain and spasm, abdominal cramping, nausea and diarrhea, hot flashes and chills, and other physiological symptoms.^2,6^ These symptoms comprise acute withdrawal, which lasts 5-7 days, or longer with long-acting opioids.^2,6,7^ Protracted withdrawal occurs after somatic symptoms subside but symptoms of negative affect persist, and can last over a year.^7^ Negative affect includes increased dysphoria, anxiety, irritability, and anhedonia.^2,6,7^ Identifying which brain regions and cell types contribute to the somatic symptoms and negative affect associated with withdrawal is critical for identifying targets for intervention. Several brain regions have been previously associated with opioid withdrawal and relapse, including the mesolimbic dopamine system, the basolateral and central amygdala, bed nucleus of the stria terminalis,locus coeruleus, raphe nucleus, the periaqueductal gray, and the habenula.^8,9^

### The habenula in addiction and withdrawal

The habenula is a well-established hub for motivation, aversion, and negative affect, with regulatory outputs to major sources of dopamine and serotonin.^10–18^ Both the medial and lateral habenula have been associated with substance use disorder and withdrawal across multiple substances,^19–24^ including nicotine,^25–27^ cocaine,^28–30^ alcohol,^31–36^ and opioids.^37^ The action of α2, α5, and β4 nicotine receptor subunits in the medial habenula (MHb) and its target the interpeduncular nucleus are both necessary and sufficient to drive nicotine withdrawal symptoms in mice.^25–27^ The habenula also regulates cocaine withdrawal and relapse, where stimulating the MHb induces reinstatement of cocaine seeking^29^ and alteration to MHb transcription factors implicated in human addiction prevent reinstatement.^28^ Cocaine withdrawal reduces GABA input to LHb neurons and restoring GABA input reduces withdrawal symptoms and reinstatement.^30^ Manipulation of LHb activity or endocannabinoid signaling in the LHb modulates self-administration of alcohol, and pharmacological interventions in the habenula can prevent depressive symptoms and hyperalgesia associated with alcohol withdrawal in mice.^31–35^

### Habenular cell types in opioid withdrawal

The habenula has an exceptionally high density of μ-opioid receptors (MORs) compared with most of the brain.^9,38–41^ MOR activation is the main driver for the positive analgesic and euphoric effects of opioids, as well as negative experiences associated with addiction and withdrawal symptoms. Complete MOR knock-out prevents morphine-induced reward, analgesia, and withdrawal in mice,^9,42^ presenting a challenge for selectively treating opioid withdrawal symptoms without antagonizing positive aspects of opioid signaling throughout the brain. A conditional knock-out of MORs in the MHb prevents aversion to the opioid antagonist naloxone and reduces somatic symptoms of withdrawal with no effect on reward or analgesia, suggesting a circuit specificity to different aspects of opioid action.^37^

The habenula is a transcriptionally diverse region with unique cell types showing a variety of anatomical projections and functional properties.^15,20,43–45^ The majority of MOR+ habenula neurons are located in the MHb and project to the interpeduncular nucleus.^39^ However, there are other MOR+ neurons in the habenula which project to the raphe nucleus and these two populations participate in different behaviors.^39^ LHb neurons projecting to the raphe nucleus exhibit withdrawal-induced synaptic depression at its inputs, mediated by cytokine signaling, which contributes to withdrawal-associated behaviors.^46^ In slice, application of DAMGO, a MOR agonist, hyperpolarizes LHb neurons and decreases excitatory input, suggesting MORs may contribute to changes in both intrinsic excitability and synaptic plasticity to control LHb firing rates.^47^ The transcriptional identity of opioid withdrawal-modulated LHb cells has not been characterized. Given that distinct cell types in the habenula have unique projection patterns with varied capability for excitatory, inhibitory, and neuropeptidergic signaling,^15,20,43–45^ understanding which of these cell types respond to opioid action is critical for building a model of how the habenula might play a role in withdrawal-associated behaviors.

Here, we identify a transcriptional marker for withdrawal-activated neurons in the habenula. Utilizing immunohistochemistry and *in situ* hybridization in spontaneous or naloxone-precipitated withdrawal, we show that glutamic acid decarboxylase 2 (GAD2) expression defines a subpopulation of LHb cells that expresses MOR and is active in opioid withdrawal. Recording fiber photometry from this population *in vivo*, we found that LHb^GAD2^ cells show a sustained increase in activity during aversive states, including acute opioid withdrawal and inescapable shock. These results establish LHb^GAD2^ cells as a target for future investigations into withdrawal and aversion circuitry in the habenula and potential intervention in the treatment of opioid withdrawal.

## Results

### Opioid withdrawal induces Fos expression in LHb neurons

To understand how opioid use and withdrawal affects the activity of habenular neurons, mice underwent a spontaneous withdrawal paradigm. Animals were given twice daily injections of either saline or an escalating dose of morphine: 20, 30, 40, 50, 60, 70, 80 mg/kg morphine s.c. for 7 days. On the 8^th^ day animals received a single 100 mg/kg morphine injection and tissue was collected 24 hours later, during spontaneous withdrawal (Figure 1A). To identify active neurons, tissue was collected and stained for the immediate early gene, Fos. There was a significant increase in Fos abundance during spontaneous withdrawal in agreement with spontaneous withdrawal from heroin.^48^ Fos^+^ cells were spatially restricted to neurons in the medial subdivision of the lateral habenula (LHbM) (Figure 1B-D) and located throughout the anterior-posterior extent of the LHbM (Figure 1C,E). Sex differences in opioid use disorder, rates of opioid overdose, and in withdrawal symptoms are well-documented in both patient populations and in animal models.^49–54^ Although there was a trend towards a sex effect wherein the increase in Fos induction appeared greater in females, statistical analysis indicated a significant drug effect (ANOVA, p< 0.01), no significant main effect on sex, and no significant interaction between drug and sex (p =0.07).

**Figure 1.**
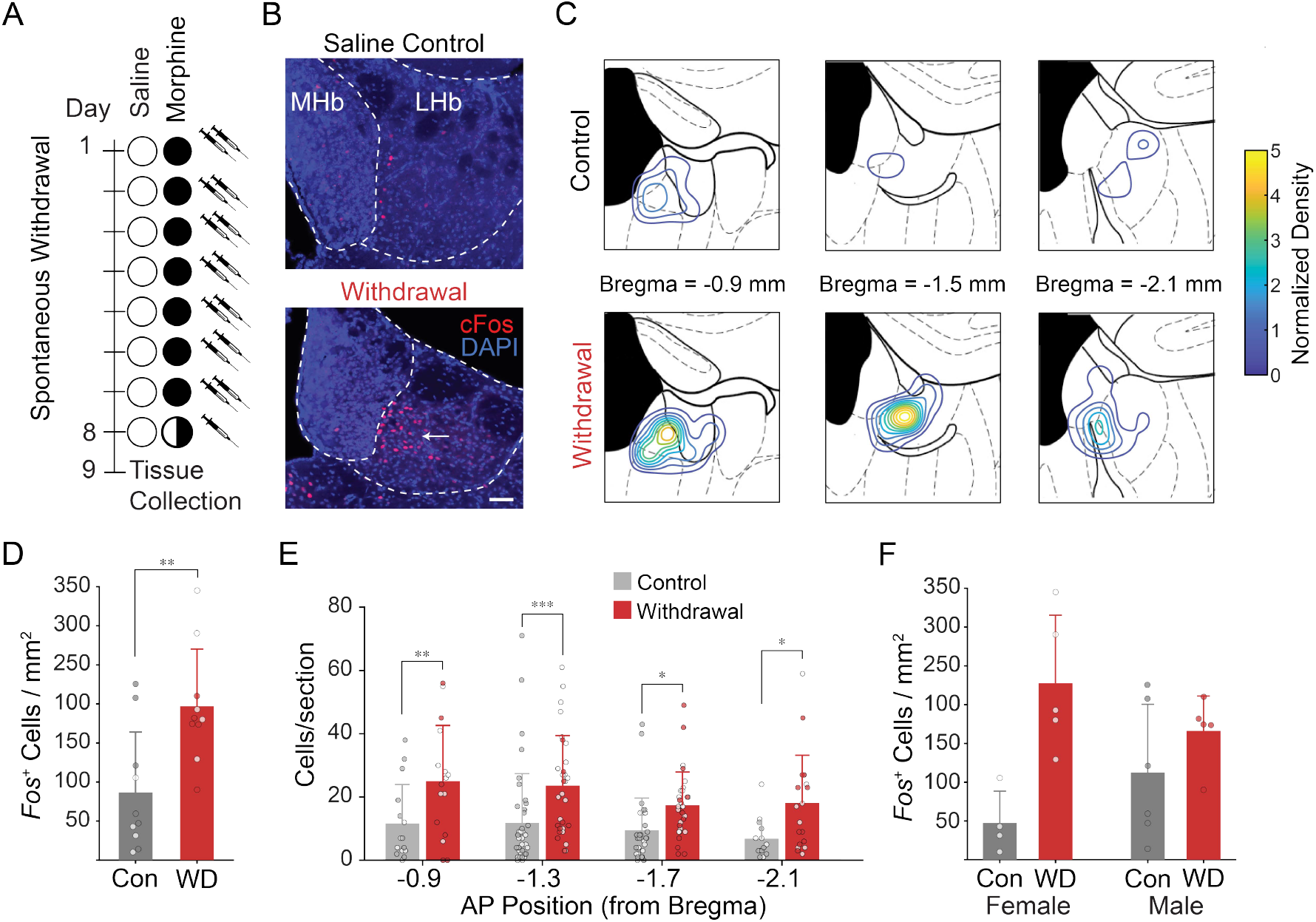
Opioid withdrawal induces Fos in the LHb. (A) Experimental paradigm for spontaneous withdrawal consisting of 7 days of twice daily escalating doses of morphine (or saline control), followed by a single final dose and 24 hours of abstinence before tissue collection. (B) Example images of anti-Fos immunostaining after spontaneous withdrawal paradigm, counterstained with DAPI. Dotted lines indicate boundaries of MHb and LHb. Arrow indicates Fos^+^ cluster in the LHb. (C) Average distribution of Fos^+^ cells across anterior-posterior axis of the habenula for each treatment condition. Contour lines indicate density of Fos^+^ cells, normalized to the maximum density in the saline condition. (D) Quantification of the density of Fos^+^ neurons in saline control and spontaneous withdrawal conditions. Each circle represents the mean for an individual animal. n=186 sections total from 10 mice per condition. **p< 0.01 by t-test. (E) Quantification of Fos^+^ cells in all sections, binned every 400 μm along the A-P axis. Each circle represents one tissue section. (F) Same data in D, separated by sex. Two-Way ANOVA, treatment effect, **p< 0.01; sex effect, n.s.; interaction, n.s. Fisher’s post hoc, **p< 0.01. Open circles indicate female, filled circles indicate male.

### Withdrawal preferentially engages LHb^GAD2^ neurons

The restricted spatial distribution of Fos expression suggested that withdrawal may recruit specific populations of LHb neurons. To identify potential cell type markers for withdrawal-activated neurons, we screened expression patterns using *in situ* hybridization data from the Allen Brain ISH Atlas^55^ and from existing data in the literature^56^ to identify genes with subregion specific expression showing a similar distribution to the *Fos*^*+*^ cells, finding that the expression of *Gad2* closely resembles the distribution of *Fos*^+^ neurons during withdrawal (Figure 2A). Using an existing data set of whole brain single cell RNA sequencing,^57^ we filtered the data set for habenular neurons and re-clustered the resulting subset.

**Figure 2.**
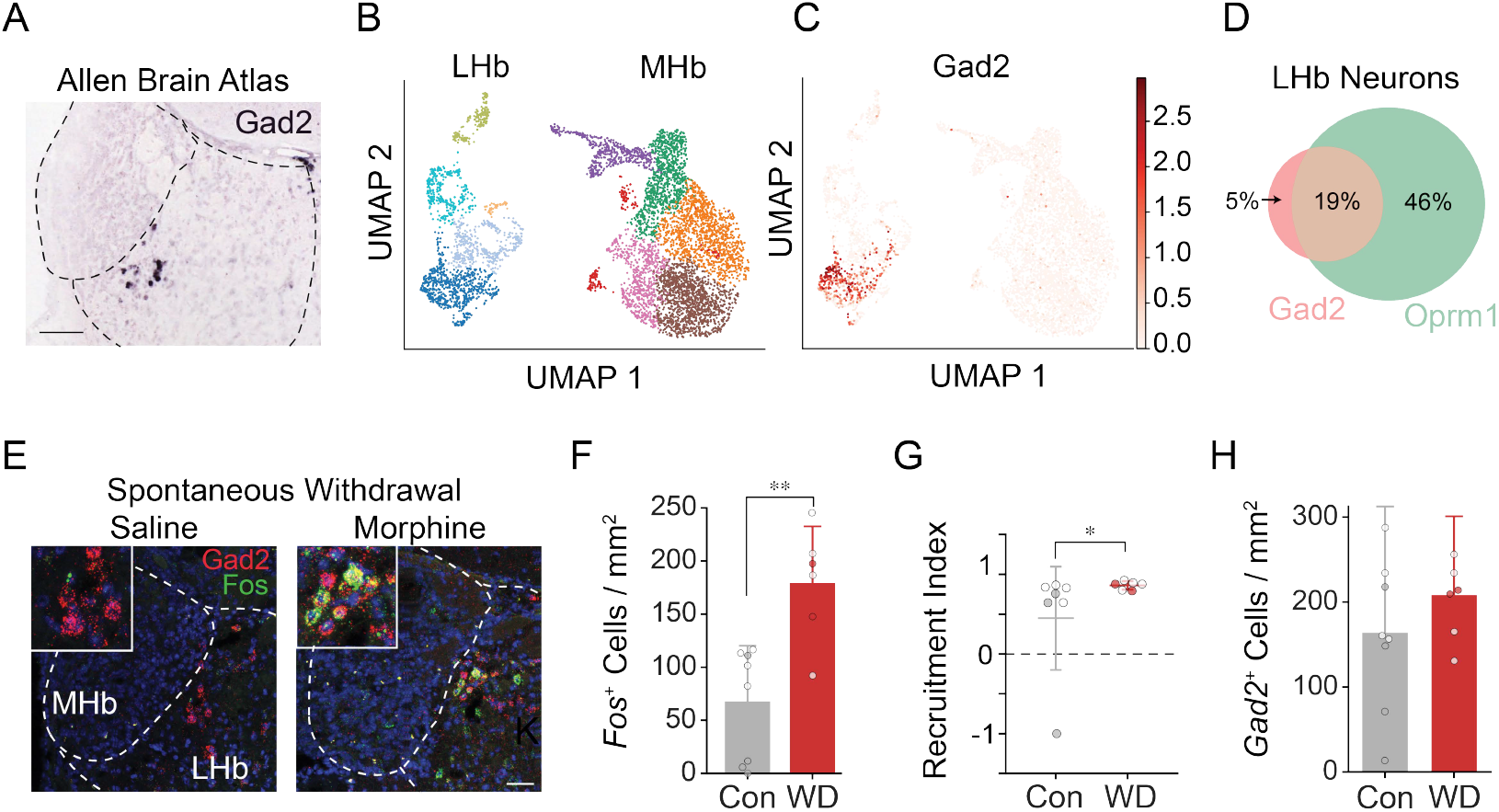
Opioid withdrawal preferentially induces Fos in LHb^GAD2^ neurons. (A) Example image of *Gad2* expression in the adult mouse habenula. Scale bar, 100 μm. Allen Mouse Brain Atlas, https://mouse.brain-map.org/experiment/show/79591669. (B) UMAP projection of single-cell RNA sequencing from Allen Institute ABC Atlas, filtered for neurons from the habenula and re-clustered using the Leiden algorithm. (C) UMAP projection in B, pseudocolored by the normalized expression of *Gad2*. (D) Overlap of *Gad2* and *Oprm1* expression in the same scRNAseq data set. Percentages indicate the proportion of LHb neurons in that condition; 30% of neurons express neither gene. (E) Example image of a double *in situ* hybridization for *Fos* and *Gad2* mRNA in the habenula during spontaneous withdrawal. Scale bar, 50 μm. (F) Mean density of *Fos*^+^ neurons in the LHb. **p< 0.01 by Mann-Whitney Test. (G) Recruitment Index, calculated the difference in probabilities of *Gad2* expression in *Fos*^+^ and *Fos*^-^ cells, divided by the sum of those probabilities. Mann-Whitney test, p<0.05. (H) Mean density of *Gad2*^*+*^ neurons in the LHb in saline control and spontaneous withdrawal. Open circles indicate female, filled circles indicate male. Error bars indicate standard deviation. n= 61 sections from 14 mice.

*Gad2* expression was well correlated with a single cluster, suggesting it might serve as a useful cell type marker (Figure 2B-c). In addition, the majority of *Gad2*^+^ cells in the data set coexpressed the μ-opioid receptor (Figure 2D). To determine if withdrawal-activated neurons express *Gad2*, we performed double *in situ* hybridization for *Fos* and *Gad2* mRNA in animals undergoing spontaneous withdrawal. In agreement with Fos immunohistochemistry, spontaneous withdrawal showed a significant increase in *Fos*^+^ cells in the LHb (Figure 2E-F). To determine if *Gad2*^*+*^ cells were overrepresented in the Fos population, a recruitment index was calculated as the difference in probabilities of *Gad2* expression in *Fos*^*+*^ and *Fos*^*-*^ cells, over the sum of those probabilities: [*p(Gad2* | *Fos*^*+*^*) - p(Gad2* | *Fos*^*-*^*)] / [p(Gad2* | *Fos*^*+*^*) + p(Gad2* | *Fos*^*-*^)]. Thus, an index of 0 indicates *Gad2*^+^ cells are equally represented in *Fos*^+^ and *Fos*^-^ populations. In the control neurons, the recruitment index was significantly greater than 0, suggesting that LHb^GAD2^ neurons are predisposed to *Fos* induction even in baseline conditions, potentially due to handling or saline control injections. However, the index was significantly increased in withdrawal relative to control with no change in the number of LHb^GAD2^ neurons (Figure 2G-H), indicating the LHb^GAD2^ neurons were preferentially recruited during withdrawal.

### LHb^GAD2^ neuron activity increases during naloxone-precipitated withdrawal

To determine how the underlying neural activity changes during withdrawal, we recorded fiber photometry from LHb^GAD2^ neurons. Interpreting the absolute amplitude of fiber photometry recordings across days is challenging, as even subtle baseline shifts due to fiber coupling placement may have a significant impact on the absolute value of the fluorescence. To allow for within session analysis of LHb^GAD2^ activity changes during withdrawal, we adopted a naloxone-precipitated withdrawal paradigm, which has been used extensively to understand the cell types and circuits contributing to withdrawal.^37,46,58–60^ To confirm that naloxone-precipitated withdrawal engages the habenular subpopulation we had identified in spontaneous withdrawal, we first analyzed the induction of *Fos* during naloxone-precipitated withdrawal. Animals received an injection of morphine (20 mg/kg) or saline control, and two hours later received naloxone dihydrate (10 mg/kg) or saline control. Tissue was collected 50 minutes after the second injection for *in situ* hybridization for *Fos* and *Gad2* mRNA. As in the spontaneous withdrawal, we saw a significant increase in the number of *Fos*^+^ neurons in the withdrawal condition (Morphine + Naloxone, Figure 3A-B), as well as significant induction of *Fos* in *Gad2*^+^ neurons (Figure 3C). There was a significant naloxone effect and morphine effect, but the interaction did not reach significance, indicating that naloxone alone may be capable of inducing Fos in LHb^GAD2^ neurons. Similarly, the recruitment index showed a significant naloxone effect, but not a significant interaction between morphine and naloxone treatments (Figure 3D). To investigate the underlying neural activity that drove *Fos* expression during naloxone-precipitated withdrawal, we recorded fiber photometry from *Gad2*^+^ neurons by performing viral injections of GCaMP8s (AAV-CAG-FLEX-jGCaMP8s-WPRE) in Gad2-Cre mice and placing a fiber optic cannula in the habenula.

**Figure 3.**
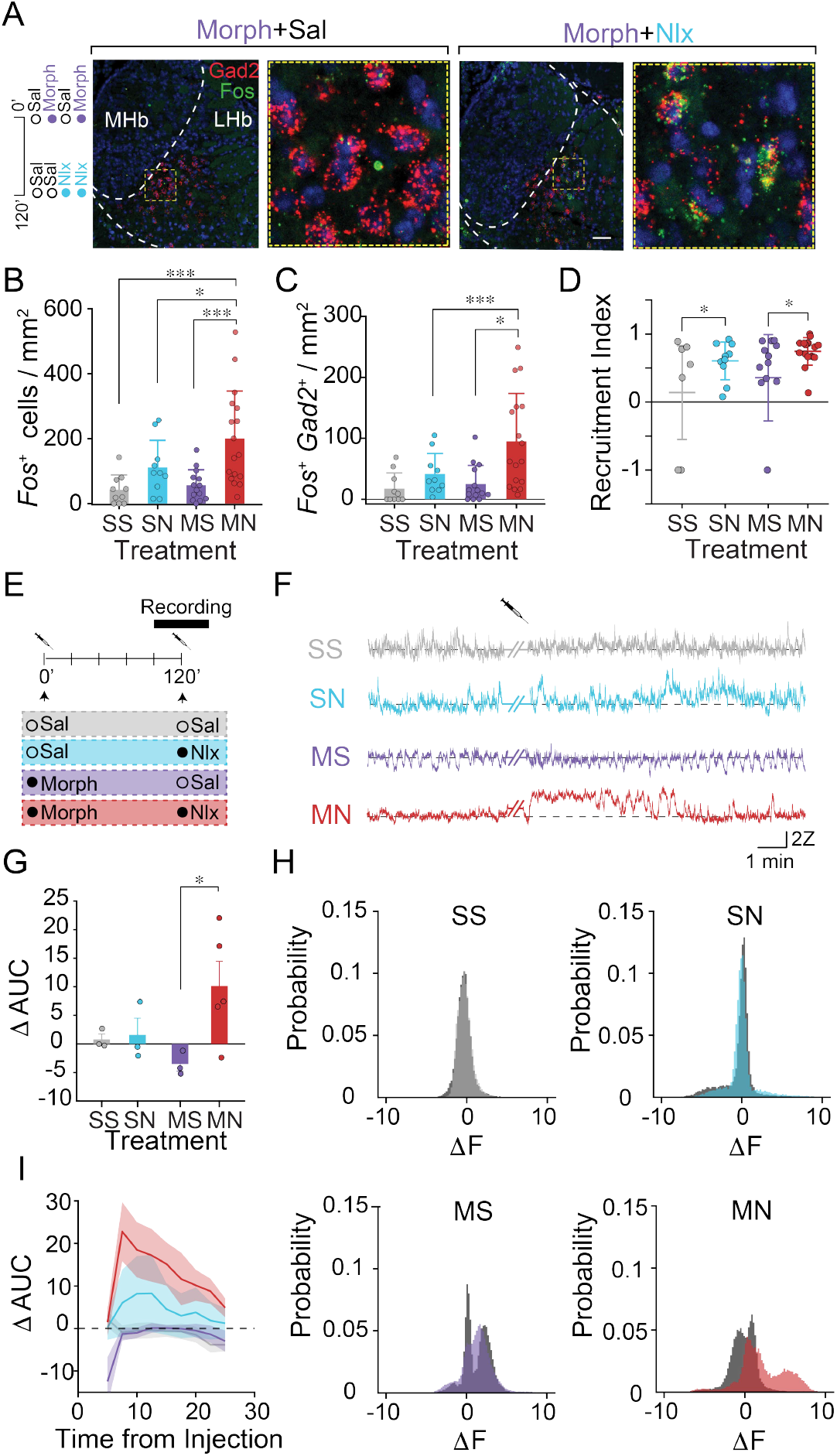
LHb^GAD2^ neurons show increased activity during withdrawal. (A) Left, experimental paradigm for acute naloxone-precipitated withdrawal. Right, example images from double *in situ* hybridization for *Fos* and *Gad2* mRNA in the habenula following naloxone-precipitated withdrawal. Scale bar, 50 μm. (B) Mean density of *Fos*^+^ neurons in the LHb. Two-Way ANOVA, Naloxone effect, p<0.001. Morphine effect, p=.07; Interaction, p=0.18. Post hoc Fisher’s test. (C) Mean density of *Fos*^+^ *Gad2*^*+*^ coexpressing neurons in the LHb. Two-Way ANOVA, Naloxone effect, p< 0.01. Morphine effect, p<0.05; Interaction, p=0.12. Post hoc Fisher’s test. (D) Recruitment Index, calculated as the difference in probabilities of *Gad2* expression in *Fos*^+^ and *Fos*^-^ populations, divided by the sum of those probabilities. ANOVA, Naloxone effect, p<0.05; post hoc Fisher’s test. (E) Experimental paradigm for recording activity from LHb^GAD2^ neurons during withdrawal. Gad2-Cre::AAV-DIO-GCaMP animals receive an injection of morphine at t = 0 min. Fiber photometry recording begins at t = 90 min. Animals are injected with naloxone or saline at 120 min and photometry recording continues to 150 min. (F) Example recordings from animals before and after the second injection of naloxone or saline. (G) Change in the area under the curve before and after naloxone or saline injection, calculated as AUC 5 min after second injection minus AUC 5 min before second injection. Dots indicate individual animals. Error bars indicate SEM. Two-Way ANOVA, Naloxone effect, *p<0.05; Morphine effect, p=0.43; Interaction, p=0.13. Post hoc Fisher’s test. (H) Histograms of the probability distribution of fiber photometry fluorescence values before (gray) and after (colored) the second injection in each treatment condition. SS, Saline + Saline; SN, Saline + Naloxone; MS, Morphine + Saline; MN, Morphine + Naloxone. n = 14 animals. Error bars indicate SEM. (I) AUC across the duration of photometry recording, calculated as a 5 min moving average of the AUC from injection onset to 25 min after the injection, relative to the AUC 5 minutes before the injection. *p<0.05, ***p<0.001

Animals were given 20 mg/kg morphine (or saline control). After 90 minutes, baseline neural activity was recorded for 30 min and then either 10 mg/kg naloxone dihydrate or saline control was administered and recording continued for 30 minutes (Figure 3E). In several morphine treated animals, LHb^GAD*2*^ neurons showed structured activity consisting of states of high and low activity (Figure 3E), as seen in the bimodal distribution of fluorescence values (Figure 3H, lower panels). Naloxone treatment resulted in a significant increase in the activity of LHb^GAD2^ neurons in animals that received morphine (Figure 3G), but not in those that received saline, and the elevated activity persisted for the duration of the recording (Figure 3I).

### LHb^GAD2^ neurons respond to aversive states

Withdrawal is an aversive state, leading to conditioned place aversion in animal models.^61,62^ To determine if LHb^GAD2^ neurons were also responsive to other aversive stimuli, we used fiber photometry in Gad2::GCaMP mice to record neural activity during the delivery of unexpected air puff. LHb^GAD2^ neurons showed increased activity in response to the air puff, though the response was modest and brief, only lasting the duration of the stimulus (Figure 4A-D). To determine if more generalized aversive states engaged LHb^GAD2^ neurons, we also recorded activity during an inescapable shock paradigm. LHb^GAD2^ neurons showed significant responses to the shock stimuli (Figure 4E-G). Unlike the response to air puff, the response to shock far outlasted the stimulus itself, only returning to baseline after ~40s (Figure 4H), suggesting that LHb^Gad2^ activity may reflect an aversive behavioral state.

**Figure 4.**
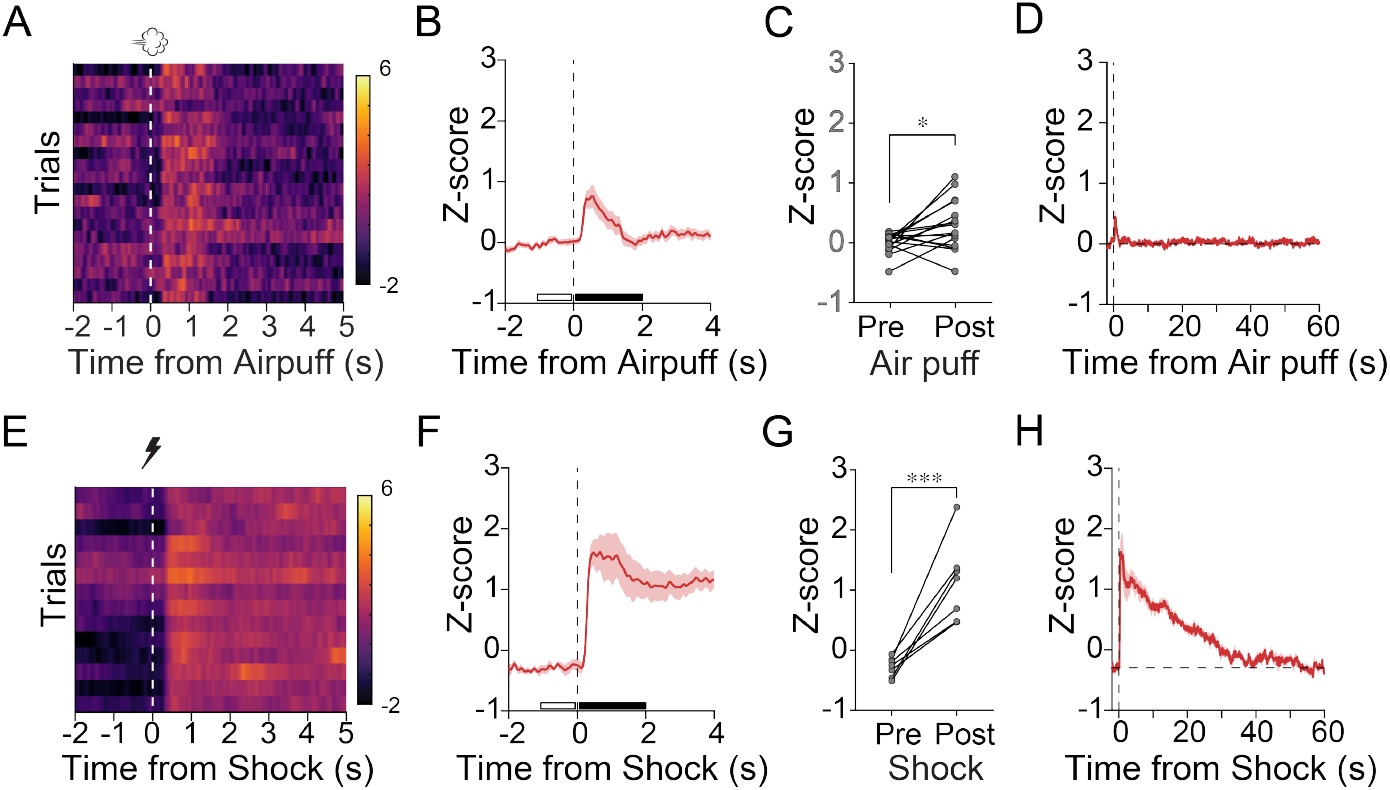
LHb^GAD2^ neurons respond to aversive stimuli and states. (A,E) Example recording of a photometry session with the presentation of (A) air puffs or (E) unpredictable shock, sorted by trial number. Colormap indicates z-score(fluorescence). (B,F) Mean z-scored fluorescence aligned to onset of 1s air puff or shock. (C,G) Quantification of change in z-score at the onset of the aversive stimulus, calculated as the mean z-score 2s after the stimulus onset minus the 1s before the onset, as indicated by the closed and open bars in B and F. Mann-Whitney test, *p<0.05, ***p<0.001. (D,H) Same data as in B,F plotted over a longer time window. Air puff, n = 16 animals; Shock, n = 7 animals. Error bars indicate SEM.

## Discussion

The habenula is associated with negative affect, aversive stimuli, and withdrawal symptoms across many substances, including opioids.^10–15,19–35,37^ The habenula is transcriptionally diverse,^43,45^ potentially providing a substrate to segregate different behavioral information. We found that a population of *GAD2*^*+*^ neurons located in the medial LHb is preferentially engaged during opioid withdrawal. *In vivo* recordings revealed a sustained increase in their activity during aversive states, including naloxone precipitated withdrawal and inescapable shock.

### Aversion and aggression

Although few studies have characterized the functional properties of LHb^GAD2^ neurons, it has been demonstrated that optogenetic stimulation or inhibition of these cells is sufficient to increase or reduce aggressive behavior respectively.^63^ Electrophysiological recordings targeted to the medial LHb, where LHb^GAD2^ cells reside, show changes to synaptic plasticity in opioid withdrawal and play a role in regulating sociability via projections to the raphe nucleus.^46^ Given that opioid withdrawal affects both sociability and aggression,^7,46,54,64^ *GAD2* expression may be a useful marker of withdrawal-activated LHbM cells, and leveraging this genetic access may allow for causal experiments to examine the potential for LHb^GAD2^ cells to serve as a modulator of withdrawal-associated behaviors, including social behaviors.

Our finding that LHb^GAD2^ cells display a sustained increase in activity during inescapable shocks and during opioid withdrawal demonstrates that LHb^GAD2^ cells carry information about aversive states, not just the stimuli themselves. Optogenetic stimulation of LHb^GAD2^ is not sufficient to generate real-time place aversion,^63^ so they may serve a modulatory role, or may drive different behavioral responses to aversion. Coupled with the evidence that stimulation promotes aggression,^63^ it is possible that the increased activity of these cells during aversive states could function to promote aggression, which may be advantageous for escaping a stressful context.

### Hormonal modulation and sex-specific differences

Although *Gad2* is used as a canonical marker for GABAergic neurons, many LHb^GAD2^ neurons coexpress glutamatergic markers. It is unclear whether they primarily form local inhibitory projections or long-range excitatory projections,^63,65^ though it has also been suggested that both types of connections may be formed by individual LHb^GAD2^ cells.^66,67^ In addition, only a subpopulation of LHb^GAD2^ cells express vesicular GABA transporter (VGAT),^67^ suggesting some may not be functionally GABAergic.^65^

LHb^GAD2^ neurons coexpressing *VGAT* also express high levels of estrogen receptors, as well as receptors for other hormones and neuromodulators.^66,68^ Gonadectomy alters both the number of *VGAT*^+^ neurons in the LHb as well as the expression levels of aromatase (the enzyme that converts androgens to estrogens) in their inputs.^66^ Because these neurons coexpress glutamatergic markers, hormonal signaling may shift the balance of individual cells between excitatory or inhibitory output.^66^ Thus, presence of neuromodulators such as estrogen may play an important role in shaping LHb^GAD2^ activity, where LHb^GAD2^ neurons could serve as a hub to integrate information about aversion with neurohormonal input to modulate behavioral response. The modulation of LHb^GAD2^ cells by sex hormones may be a potential mechanism for the trend we saw towards sex differences in Fos induction in this circuit (Figure 1F). Both sexes were included in our photometry data; while animals of both sexes showed prominent increases in activity during withdrawal, our data were underpowered to make definitive claims about sex differences.

### LHb^GAD2^ circuit connectivity

We show that LHb^GAD2^ neurons coexpress MORs (Figure 2D), but it is unclear if the change in LHb^GAD2^ activity is due to direct modulation of LHb^GAD2^ excitability, or from changes in the activity of their inputs. Partial knockout of MORs in the habenula is sufficient to reduce somatic symptoms of opioid withdrawal.^37^ However, the knockout targeted the MHb, where MOR density is highest,^37,41^ likely sparing LHb^GAD2^ neurons. It has been shown that Hb MOR+ cells that project to the interpeduncular nucleus and raphe nucleus participate in different behaviors,^39^ suggesting that MHb and LHb MOR+ cells, which preferentially target these regions respectively, could participate in different functions. We found no increase in Fos for MHb neurons, so any withdrawal-associated changes in MHb MORs are likely not due to changes in their firing rates, but we cannot exclude potential modulation of synaptic release by axonally localized MORs.

Changes in LHb^GAD2^ cell activity may occur due to changes in intrinsic excitability or due to changes in their inputs, potentially inputs arising from the lateral hypothalamus. Stimulating lateral hypothalamus projections to LHb^GAD2^ cells, or altering their ability to carry out orexinergic signaling to LHb^GAD2^ cells, is sufficient to drive changes in LHb^GAD2^ cell activity and behavior.^63^ Orexin^+^ cells in the LH also express MORs and tend to increase both Fos and Orexin production following opioid withdrawal.^59,69^ The locus of MOR action may also depend on the withdrawal paradigm. We show both spontaneous and naloxone-precipitated withdrawal increases Fos labeling, but spontaneous withdrawal more specifically recruits LHb^GAD2^ neurons. Chronic exposure may engage long-term plasticity mechanisms where changes in LH →LHb^GAD2^ synaptic transmission contribute to withdrawal-induced activity and confer specificity to LHb^GAD2^ neurons.

Our findings identify a transcriptional marker of a LHb cell population that increases activity in response to aversive stimuli and opioid withdrawal. There has been increasing interest in this LHb^GAD2^ population due to their role in aggressive behavior, local and distant projection patterns, and unique transcriptional profile.^63,65–67^ Our work suggests that LHb^GAD2^ cells may have unique translational relevance for processing aversive stimuli and states, including opioid withdrawal.

## Methods

### Mice

Animal husbandry and all aspects of animal care and euthanasia as described were in accordance with guidelines from the NIH and have been approved by members of the University of Oregon Institutional Animal Care and Use Committee. Male and female mice aged 10-20 weeks were used for all experiments. Animals were group housed whenever possible. All mice were either C57Bl/6J (Jax#000664) or Gad2-Cre (Gad2^tm2(cre)Zjh^/J, Jax#010802).

### Naloxone-Precipitated Withdrawal

Animals were acclimated to handling and injections for 3 days using a once-daily saline injection. At t=0, animals were either injected with morphine (20 mg/kg, s.c.) or an equivalent volume of saline. After 2 hours, animals were injected with either naloxone (10 mg/kg, s.c.) or an equivalent volume of saline. For Fos immunohistochemistry experiments, animals were perfused 110 minutes after the second injection. For *in situ* hybridization, animals were perfused 50 minutes after the second injection.

### Spontaneous Withdrawal

Animals received twice-daily escalating doses of morphine for 8 days (in mg/kg): 20, 30, 40, 50, 60, 70, 80, 100. Injections were separated by 12 hours, typically zeitgeber (ZT) time 1hr and 13hr. On day 8, only the ZT1 injection was administered. After 24 hours of abstinence (or control), animals were perfused for immunohistochemistry or *in situ* hybridization.

### Air puff

Animals were placed in a chamber (18 cm x 18 cm x 30 cm) with a metal grated floor and allowed to explore freely. An air diffuser box was positioned below the chamber floor. Air puffs covering the full field of the chamber were delivered using house air (60 psi) for 1 s using a pneumatic valve controlled by a Bpod behavioral control system (Sanworks). Trials were separated by a variable inter-trial interval (μ = 30 s, σ = 5 s). Air puff trials and sham trials in which a dummy valve was opened were presented with equal likelihood and pseudo-randomly interleaved in the session. Animals were exposed to 20 air puffs during a single session.

### Foot shock

Animals were placed in a chamber (18 cm × 18 cm × 30 cm) with a metal grated floor. Foot shocks were delivered to the grid floor by a precision animal shocker (Lafayette Instrument, Model HSCK100AP). Foot shocks were controlled by an Arduino-based behavioral control protocol (BehaviorDEPOT^70^). Sessions started with a 30 s habituation period preceding the first trial. For each trial, a mild foot shock (0.1 mA) was administered for 1 s. Each shock presentation was followed by a variable inter-trial interval of 60 – 120 s. Animals were exposed to a single session with a total of 20 shocks.

### Immunohistochemistry

To prepare tissue for immunostaining, mice were anesthetized with Euthasol (0.5 ml/kg delivered IP, 390 mg pentobarbital sodium (barbituric acid derivative), 50 mg phenytoin sodium, MWI Animal Health) and transcardially perfused with fixative containing 4% PFA (Electron Microscopy Sciences, 1224SK). Brain tissue was removed from the skull and post-fixed overnight at 4°C. Tissue was sectioned with a vibratome (TPI 1000) at a thickness of 50-75 μm. Tissue was washed 3 times (15 min each) in 1xPBST and incubated in blocking buffer (1x PBS, 0.01% TX-100, 10% Normal Donkey Serum (Jackson ImmunoResearch 017-000-121)) for 30 minutes. Primary antibodies were diluted in blocking buffer and tissue was incubated overnight at 4°C. The primary antibody dilutions are as follows: rabbit anti-Fos (Santa Cruz Biotech, sc-271243 at 1:1000) or mouse anti-Fos (Synaptic Systems, #226003 at 1:1000). Tissue was washed three times for 15 minutes each and then incubated in a secondary antibody solution diluted in blocking buffer for 1 hour at room temperature. Secondary antibodies were donkey-anti-rabbit-647 (Jackson ImmunoResearch, 711-605-152) and goat-anti-mouse-647 (Thermofisher, A-21235). Tissue was washed three times (5 minutes each), mounted on slides, and coverslipped with FluoromountG with DAPI (Fisher Scientific, 50-112-8966).

### In situ hybridization

To prepare tissue for *in situ* hybridization, mice were anesthetized with isoflurane and rapidly decapitated. Brain tissue was immediately removed, embedded in OCT, and flash frozen in liquid nitrogen. Tissue was equilibrated to -20°C and sectioned on a microtome (Leica CM3050 S) to a thickness of 20 μm and mounted on Superfrost Plus slides (Fisher Scientific, 12-550-15). Tissue was promptly fixed in 4% PFA (Electron Microscopy Sciences, 1224SK) for 10 minutes, then transferred to -20°C methanol (Sigma-Aldrich, 179337) and incubated at -80°C for 1 hour. Tissue was washed two times in 5xSSC, then incubated in probe hybridization buffer (Molecular Instruments) for 30 minutes. Probes targeting *Fos* and *Gad2* mRNA were designed by Molecular Instruments and diluted to 2 nM final concentration in hybridization buffer. Hybridization buffer with probes was added to the slides, covered with Hybrislips (Fisher Scientific, H18202), and incubated overnight in a humidified chamber at 37°C. Slides were washed with probe wash buffer (Molecular Instruments) and 5xSSCT, then equilibrated in amplification buffer (Molecular Instruments) for 30 min. Fluorophore-labeled hairpins (Molecular Instruments) were heated to 95°C for 90 seconds, then moved to room temperature for 30 min to cool. Cooled hairpins were added to amplification buffer and the resulting solution was added to tissue sections and coverslipped. The amplification reaction was run overnight at room temperature, protected from light exposure. Amplified sections were washed 4 times with 5xSSCT in a Coplin jar and then coverslipped with FluoromountG with DAPI for confocal imaging.

### Confocal Microscopy and Image Analysis

Images from immunohistochemistry and *in situ* hybridization assays were acquired on a Zeiss laser scanning confocal microscope (LSM700). For immunohistochemistry, images were acquired with a 20x objective at 2x digital zoom. Six z-planes were taken with a 3 μm interval and max-projected for analysis. For *in situ* hybridization, images were acquired with a 20x objective at 2x digital zoom. Four z-planes were taken with a 3 μm interval and max-projected for analysis. Confocal Images were analyzed in custom MATLAB code. LHb boundaries were drawn using the DAPI channel. Fos-expressing cells were manually identified in each section. If additional channels were acquired (Gad2), each Fos-expressing neuron was manually assessed for coexpression. Cell densities were calculated using the LHb boundaries and the number of cells for each condition. To generate cell contours to depict mean cell densities across multiple animals and sections, a geometric transformation was performed between an Atlas Image (Paxinos and Franklin, 2001) and the DAPI image for each section. The transformation was applied to all cell locations and then binned (10 μm x 10 μm). Cell densities in each bin were normalized to the maximum bin density for control (saline) treatment and contour lines were plotted for each treatment. Recruitment Index was calculated by difference over the sum: [*p(Gad2* | *Fos*^*+*^*) - p(Gad2* | *Fos*^*-*^*)] / [p(Gad2* | *Fos*^*+*^*) + p(Gad2* | *Fos*^*-*^)]. Values are bounded by -1 and +1. An index of 1 indicates that all *Fos*+ cells are *Gad2*^+^ and no *Fos*^−^ cells are *Gad2*^+^, reflecting maximal specificity. An index of −1 indicates the opposite extreme, where *Gad2*^+^ cells are found exclusively among *Fos*^−^ neurons, reflecting maximal exclusion from the active population.

### Fiber photometry

Surgical Procedures: Male and female mice age 11-18 weeks were anesthetized with 4% isoflurane and maintained at 1-2% isoflurane mixed with oxygen at a flow rate of 1.5 – 2 L/min. Depth of anesthesia was monitored throughout surgery using tail pinch responses and breathing rates. Mice were mounted on a stereotaxic surgical station (David Kopf Instruments). A small craniotomy was made over the lateral habenula and stereotaxic injections were used to deliver viruses. 300 nL of AAV-CAG-FLEX-jGCaMP8s-WPRE was injected into the right LHb (AP: -1.4 mm, ML: -0.5 mm, DV: -3.2 mm) at a rate of 30 nl/min and a fiber optic cannula (Doric Lenses, MFC_400/430-0.66_3.5mm_MF2.5_A30 or MFC_400/430-0.66_3.5mm_MF2.5_TP1.0) was placed above the injection site (AP: - 1.4 mm, ML: -0.5 mm, DV: -3.15 mm). Coordinates are relative to bregma, with DV coordinates relative to skull surface. Cannulas were cemented in place with Metabond and Amalgambond Catalyst (Parkell) and animals were allowed to recover for at least 2 weeks prior to handling and data recording.

Recording: Fiber photometry signals were recorded with a fiber optic patch cord (Doric Lenses, MFP_400/430/1100-0.57_1m_FC-MF2.5_LAF) coupled to the animals with a ceramic or brass sleeve. GCaMP was excited with LEDs emitting 405 nm or 465 nm light, modulated with a lock-in amplifier (RZ10X LUX-I/O system, Tucker Davis Technologies). LED light sources were sinusoidally modulated at 210 and 330 Hz, respectively, and filtered through a fluorescence minicube (Doric Lenses, FMC6_IE(400-410)_E1(460-490)_F1(500-540)_E2(555-570)_F2(580-680)_S). Fluorescence was captured by integrated photosensors and processed with a RZ10X LUX-I/O digital acquisition system and Synapse software (Tucker-Davis Technologies). Photometry data were synced to behavioral data using TTL outputs captured by Synapse software to sync with injection timing, air puffs, and shocks. For each behavioral session, the 405 nm signal was fit to the GCaMP signal and subtracted from it. The resulting signal was downsampled to 20 Hz and reported as Normalized F. Z-scores were calculated using the Normalized F across the entire behavioral session. Comparisons across animals and genotypes were reported as z-score to reduce effects of variability in viral expression across animals.

Histology: Following all experiments for each cohort, animals were deeply anesthetized using an intraperitoneal injection of Euthasol (0.5 ml/kg delivered IP, 390 mg pentobarbital sodium (barbituric acid derivative), 50 mg phenytoin sodium, MWI Animal Health), transcardially perfused with phosphate-buffered saline (PBS) and fixative containing 4% PFA (Electron Microscopy Sciences, 1224SK), and postfixed 48 hours at 4 °C. Brains were sectioned at 50 μm thickness near the site of cannula placements. Brains were permeabilized in 1xPBS with 0.1% TX100 and incubated overnight in anti-GFP antibody solution (Thermo Fisher Cat # A-31852 or Cat # A-21311 at 1:2000 dilution in 1xPBS with 1% Bovine Serum Albumin) to stain GCaMP-expressing neurons. Sections were washed in 1xPBS and mounted on coverslips with Fluoromount-G with DAPI (Fisher Scientific, 50-112-8966). Brain sections were imaged on a Nikon ‘Sora’ Spinning Disk Confocal Microscope with 405 nm and 640 nm lasers and a 20x objective.

## Resource Availability

### Lead Contact

Requests for further information and resources should be directed to and will be fulfilled by the lead contact, Emily Sylwestrak.

### Materials Availability

This study did not generate new materials.

### Data and Code Availability

- All data generated in this study will be shared by the lead contact upon request.
- This paper does not report original code.
- Any additional information required to reanalyze the data reported in this paper is available from the lead contact (ES) upon request

## Acknowledgements

We thank the entire Sylwestrak Laboratory for helpful discussions. We also thank the University of Oregon Terrestrial Animal Care Services staff for the diligent care of our animals, and Adam Fries of the Genomics and Cell Characterization Core for technical imaging support. This work was funded with the generous support of the Brain Research Foundation (BRFSG-2020-08).

## Author Contributions

L.B. contributed to conceptualizing experiments, formal analysis, project administration, and writing of the original manuscript, and conducted immunohistochemical experiments, and fiber photometry recordings. V.M., B.H., and I.S. conducted immunohistochemistry experiments. E.L.S contributed to supervision, conceptualizing experiments, data curation, formal analysis, funding acquisition, project administration, resources, and writing of the original manuscript.

